# PI4K2β constrains non-canonical Wnt-PCP signalling and is localised by PAR-1 (MARK2/3) phosphorylation

**DOI:** 10.1101/660431

**Authors:** S. M. Tsang, William Cheng, Jingjing Li, Jeremy B. A. Green

**Affiliations:** Centre for Craniofacial and Regenerative Biology, King’s College London, Guy’s Tower Floor 27, London SE1 9RT, UK

## Abstract

Canonical Wnt signalling is critically important in embryonic cell-type specification and cancer, while non-canonical Wnt signalling is primarily implicated in physical morphogenesis, especially planar cell polarity (PCP). Both are modulated by the polarity kinase PAR-1 (MARK2/3). PAR-1 phosphorylates the Wnt transducer Dishevelled, but there is evidence that it exerts control through other targets. Here we describe an *in vitro* screen for new targets of PAR-1 in which we identified phosphatidyl-inositol-4-kinase-2-beta (PI4K2β) as a substrate. Perturbation phenotypes and reporter assays *in vivo* show that PI4K2β inhibits both canonical and non-canonical Wnt pathways, in contrast to PI4K2α, which promotes canonical but does not affect non-canonical signalling. We show that PI4K2β acts in Wnt-responding tissue, not in Wnt production or secretion. Subcellularly, PI4K2β is cortically enriched, unlike PI4K2α, and is basolateral in polarised cells. Mutation of the PAR-1 phosphorylation site of PI4K2β mis-localises it and the endogenous core PCP protein, Vangl2. Our results reveal that PAR-1 interacts with the vertebrate PCP signalling pathway via PI4K2β.

## INTRODUCTION

The polarity protein partitioning-defective 1 (PAR-1), also known as microtubule affinity regulating kinases (MARK2 or −3), is a highly conserved serine/threonine kinase with a number of polarity-associated functions ^1^. PAR-1 plays essential roles in asymmetric cell division and the establishment of cell polarity from invertebrates to mammals ^2^. PAR-1 interacts with multiple signalling pathways including the canonical and non-canonical Wnt signalling pathways ^3, 4, 5, 6^. However, regulation and cross-talk between PAR-1 and Wnt signalling components are not well understood. Both canonical and non-canonical Wnt pathways are initiated by binding extracellular Wnt protein, which acts through a Frizzled (Fz) receptor and Dishevelled (Dvl) to activate downstream targets. Canonical Wnt pathway leads to stabilisation/translocation of β-catenin and transcriptional activation of target genes ^7^, while non-canonical Wnt pathways diverge at the level of Dvl into the Planar Cell Polarity (PCP) pathway involving Rho/Rac small GTPase and Jun N-terminal kinase (JNK) or the Wnt/Ca^2+^ pathway via protein kinase C (PKC) ^8^. Functionally, the canonical pathway specifies dorsal structures during early *Xenopus* development ^9^, while the non-canonical pathways regulate cell polarity including that required for convergent extension movements during zebrafish and *Xenopus* gastrulation ^10, 11^.

The PCP pathway promotes polarisation in the plane of the epithelial cells, perpendicular to their apicobasal axis ^12^, affecting cell morphology, at least in part independently of transcription ^13^. A key feature of this pathway is the asymmetric distribution of their core PCP components such as transmembrane proteins Frizzled (Fz) and Van Gogh (Vang) and cytoplasmic regulators Dishevelled (Dvl) and Prickle (Pk) both within and between cells ^14^. Disruption of the core PCP proteins is associated with developmental abnormalities such as defects in convergent extension and neural tube closure ^15, 16^ and also implicated in cancer invasiveness ^17^. Van Gogh-like (Vangl) is a mammalian homologue of the Drosophila PCP gene Van Gogh (Vang) (also known as Strabismus). Vang/Vangl is known to be involved in the non-canonical PCP pathway via the activation of JNK ^18^. Studies have shown Vang/Vangl binds directly to Dvl ^18^ and Prickle ^19^ and is involved in their recruitment to the plasma membrane. The molecular networks that underlie its membrane targeting however, remain largely unknown.

Phosphatidylinositol (PI) can be phosphorylated at the 3-, 4- and 5-positions of the inositol ring by different lipid kinases to generate seven distinct phosphoinositides that mediate important signalling and trafficking functions ^20, 21^. Phosphatidylinositol 4-kinases (PI4Ks) generate phosphatidylinositol 4-phosphate (PI4P) and are structurally and biochemically classified into type II (PI4K2) and type III (PI4K3) families ^20^. Phosphatidylinositol 4-kinase type 2 (PI4K2) is the predominant PI4-kinase activity in cell extracts. Two structurally related isoforms, PI4K2*α* and PI4K2*β*, with similar enzymatic properties contribute to this activity.

Despite similarities between PI4K2α and -β, accumulating evidence suggests that they have non-identical biological functions. In cultured mammalian cells, PI4K2α is catalytically active, tightly membrane-associated, and predominantly located to intracellular membranes such as *trans*-Golgi network ^22, 23^ and early and late endosomes ^24, 25, 26^. PI4K2α plays essential roles in endocytosis ^22^, Golgi function ^24^ and trafficking processes such as the delivery of AP-3–dependent cargoes to lysosomal compartments ^25, 27^ and lysosomal degradation of epidermal growth factor receptors ^26^. Deletion of PI4K2α gene in mice causes late onset neurodegeneration ^28^. In addition, PI4K2α promotes LRP6 phosphorylation during canonical Wnt signalling in both mammalian cells and *Xenopus* ^29, 30^. Much less is known about PI4K2β. Like PI4K2*α* PI4K2β localises to membranes of the Golgi and endosomal systems ^24 23^ and lacks a transmembrane domain but associates tightly with membranes through multiple palmitoylation sites ^23 31^. However, PI4K2α and PI4K2β differ in their membrane recruitment: PI4K2*α* is constitutively membrane associated, whereas PI4K2*β* is dynamically palmitoylated and recruited to the plasma membrane in response to platelet-derived growth factor stimulation or Rac activation and in turn becomes activated ^23^. PI4K2*β* therefore appears to be regulated by membrane recruitment. PI4K2β does not activate canonical Wnt signalling ^29^ but has been associated with inhibition of invasiveness of tumour cells^32^.

Previously, we showed that different PAR-1 isoforms are involved in *Xenopus* development and that each is associated with a different branch of Wnt signalling pathway ^6^. In most vertebrate species examined, the isoforms are splice variants. Isoform PAR-1BY (MARK2 variant Y) controls the localisation of one of its substrates, Dishevelled (Dvl), in the non-canonical Wnt planar cell polarity (PCP) signalling pathway. In contrast, isoforms PAR-1A (MARK3) and PAR-1BX (MARK2 variant X) primarily act on canonical signalling to β-catenin. Interestingly, it was also found that a PAR-1-non-phoshorylateable mutant of Dvl was able to induce a canonical Wnt signal just as potently as wild type Dvl ^6^. This suggests that phosphorylation of Dvl may not be the only target of PAR-1 in Wnt-β-catenin signal transduction.

Here we describe an unbiased screen for novel phosphorylation targets of PAR-1 in which we identified PI4K2β as a potential target. We then analysed the developmental role of PI4K2β and found that depletion produces phenotypes consistent with roles in inhibiting both canonical and non-canonical Wnt signalling. These were confirmed with reporter assays, which were further used to show that the effects on canonical signalling were in the responding rather than the Wnt-secreting cells. We also observed that PI4K2β is basolaterally localised in polarised epithelial cells in *Xenopus*, and that mutation of the PAR-1 phosphorylation site is sufficient to abrogate such localisation. Furthermore, we showed PI4K2β co-localises with a key PCP protein Vangl2 at the basolateral domain and Vangl2 is mis-localised by expression of the PAR-1-unphosphorylatable PI4K2β mutant protein. Our data suggests that PAR-1-dependent PI4K2β basolateral localisation plays an important role in the regulation of the vertebrate PCP signalling pathway.

## RESULTS

### Phosphatidylinositol 4-kinase type II beta (PI4K2β) is a novel phosphorylation target of PAR-1

We previously showed that a PAR-1-non-phoshorylatable mutant of Dvl was fully able to induce a canonical Wnt signal, and that PAR-1A could super-induce signalling triggered by this mutant ^6^. This suggests that PAR-1 kinase could act by potentiating canonical signalling to Wnt-β-catenin via alternative (non-Dvl) target(s). We therefore carried out an unbiased screen for novel phosphorylation targets of PAR-1 to understand its mechanism of action in Wnt signalling branches. To screen for novel phosphorylation targets of PAR-1, pools of clones from a 7000-clone arrayed normalized *Xenopus laevis* oocyte cDNA library ^33^ were first transcribed and translated *in vitro* in the presence of non-radioactively labelled amino acids. The resulting protein pools were then incubated with purified recombinant GST-tagged human MARK3 wild type (WT) or kinase dead (KD) proteins ^34^ and subsequently analyzed by SDS-PAGE and Western blotting (Fig.1A). Proteins phosphorylated after kinase treatment were identified by looking for alterations in their mobility in gel electrophoresis. “Band shifted” clones identified in the pool screen were further confirmed individually to verify the result. Using this approach, we identified 14 potential targets and 11 of them were known genes or close homologues (Fig.S1A). Among them, we selected phosphatidylinositol 4-kinase type II beta (PI4K2β) for further investigation. PI4K2β is an inositide kinase that participates in the phosphatidylinositol cycle. Western blot analysis in Fig.1B shows that PI4K2β when tested individually becomes phosphorylated after incubation with MARK3 wild-type kinase and migrated with an increased mobility upon gel electrophoresis compared to the kinase dead-treated control. Although phosphorylation increases the molecular weight of a protein, an increase in mobility is not unprecedented (e.g. upon T-loop phosphorylation of CDK1 ^35^) and presumably results from secondary structural changes, or phosphorylation-facilitated modifications in the cell free extract, that facilitate migration through the pores of the electrophoresis gel. It is known that both *Xenopus* PI4K2α and-β contain the known PAR-1 phosphorylation consensus site KVGS at Ser-255 and 265 respectively (Fig.S1B) but neither has previously been identified as a PAR-1 substrate.

**Fig.1.**
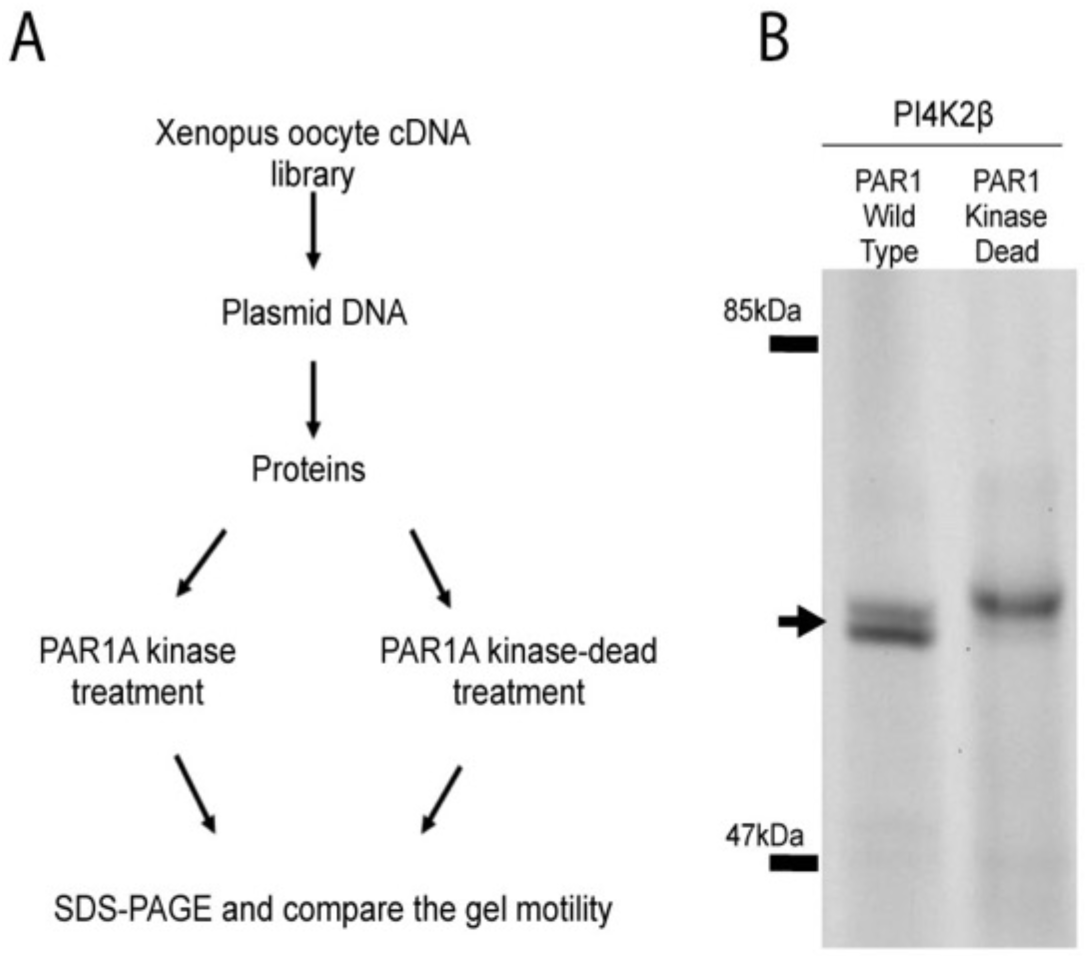
Phosphatidylinositol 4-kinase type II beta (PI4K2β) as a novel phosphorylation target of PAR-1. A) Schematic diagram of the *Xenopus* expression library screen. A 7000-clone arrayed normalized *Xenopus laevis* oocyte cDNA library was used. B) Western blot analysis showing that PI4K2β *in vitro*-transcribed and translated from cloned DNA becomes phosphorylated after incubation with MARK3 wild-type (WT) kinase and migrated with an increased mobility upon gel electrophoresis as compared to the MARK3 kinase dead (KD)-treated protein. Purified GST-tagged human MARK3 wild type (WT) and kinase dead (KD) proteins were used. Numbers at the left are molecular mass in kDa. Arrow indicates the PI4K2β protein becomes phosphorylated with a reduced mobility after kinase treatment.

### The *in vivo* function of PI4K2β *during Xenopus* embryogenesis

PI4K2α was previously implicated in Wnt signalling as an important cofactor in canonical signalling ^29^, but PI4K2β has not previously been analysed *in vivo*. To evaluate the *in vivo* function of PI4K2β during *Xenopus* development, we first examined whether *Xenopus* PI4k2β is expressed during gastrulation. Analysis of the spatial expression of the PI4k2β transcripts during *Xenopus* early embryogenesis by whole-mount *in situ* hybridization revealed ubiquitous expression of the transcripts (data not shown).

Next, we carried out the loss-of-function analysis by using translation-blocking morpholino oligonucleotides (Mos) against its mRNA at the 5’-UTR. Two non-overlapping Mos (designated PI4K2β-Mo-1 and PI4K2β-Mo-2) had the same effects as follows. Embryos injected with 15ng of either PI4K2β-Mo into two dorsal blastomeres at 4-cell stage had no discernable effect up to gastrulation (data not shown). However, at tailbud and tadpole stages, more than 60% of the PI4K2β-depleted embryos (n=51) displayed severe defects in dorsoanterior structure (Fig.2A-F&M, Fig.S2A). The phenotype was variable but dose-dependent and an apparent combination of two often distinct defects: reduced or missing head and a severely dorsally bent body axis. Absence of anterior structures if accompanied by loss of notochord can, in principle, be due to partial loss of the Spemann Organiser. Examination of the PI4K2β-depleted embryos showed the presence of well-differentiated, albeit disorganised notochord tissue (Fig.2F). This implicates an excess of late posteriorising canonical Wnt signalling which has previously been associated with loss of anterior structures with retention of notochord ^36^. Notochord disorganisation suggested a defects in the control of convergent extension movements, which are regulated by non-canonical Wnt-planar cell polarity (PCP) signalling ^37, 38^. Neither ventral injections (not shown) nor control Mo (Fig.2A-C) had any overt effects.

**Fig.2.**
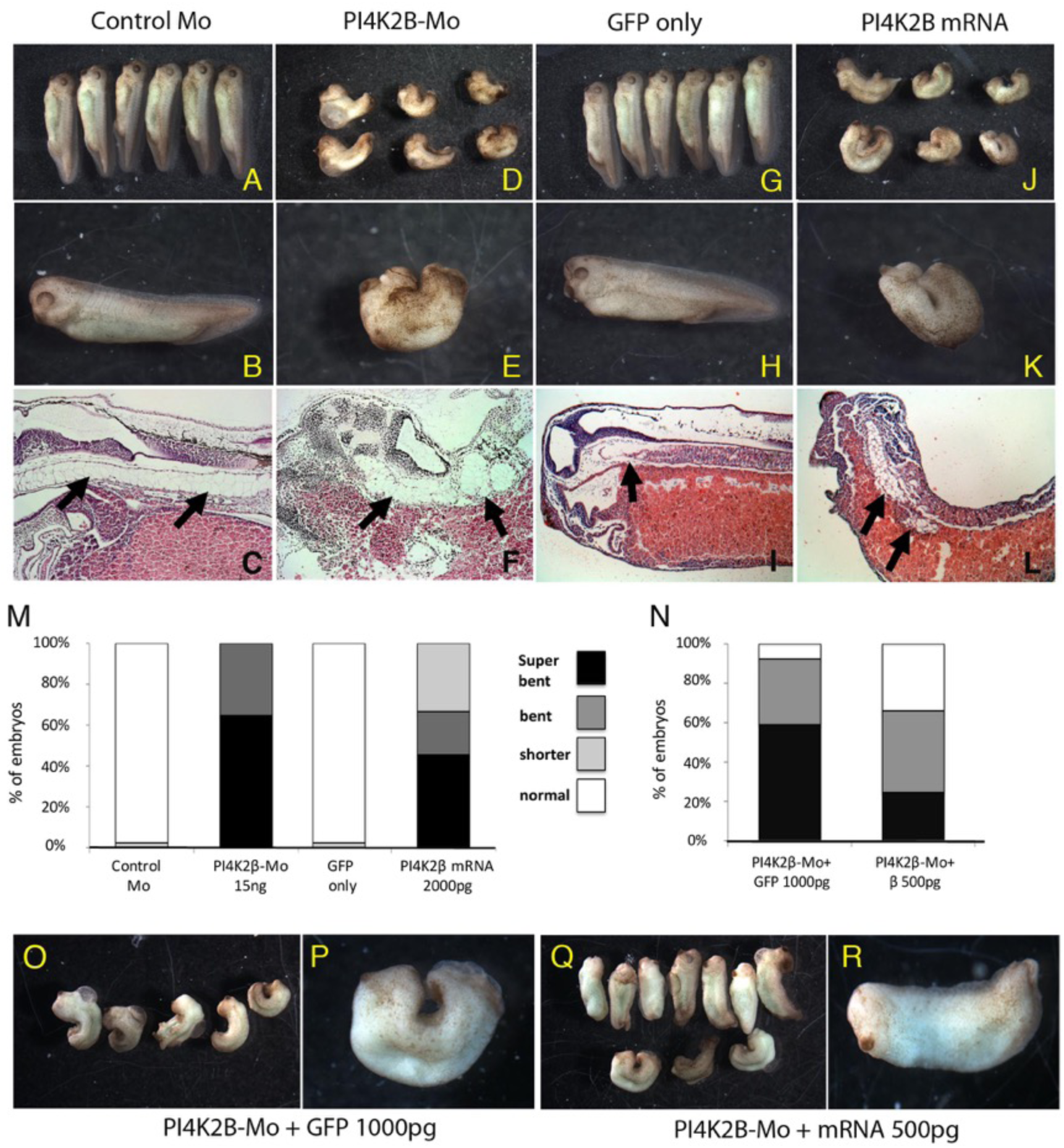
Both gain- and loss-of-function of PI4K2β cause severe defects in dorsoanterior structure during embryogenesis. (A-L) Representative tailbud stage embryos injected as indicated at 2-4 cell stage with control Mo 15 ng or GFP mRNA 2000 pg versus PI4K2β-Mo 15 ng or mRNA 2000 pg, show dorsally bent axes in PI4K2β-perturbed cases D, E, J, and K. Hematoxylin-Eosin-stained sagittal sections (C,F,I,L) show the presence of notochord in all cases (black arrows) but disorganised in PI4K2β-perturbed embryos (F,L). (M&N) Histograms quantifying frequencies of the phenotypes. Data summarise at least three independent experiments with at least 20 embryos in each condition for each. (N -R) Rescue experiments by co-injecting Mo with GFP- or full length PI4K2β mRNA. 500pg of PI4K2β mRNA, but not GFP alone, partially rescues the convergent extension defect to give a shorter but straight axis (N,Q-R).

For gain-of-function analysis, we injected PI4K2β mRNA into two dorsal blastomeres at 4-cell stage. The embryos injected with up to 2000pg PI4K2β mRNA also resulted in a shorter and dorsally bent axis (Fig.2J-L). As shown in Fig.2M, more than 50% of the PI4K2β mRNA-injected embryos (n=50) have the similar “super bent” phenotype to that of embryos injected with PI4K2β-Mo. H&E stained sections of the PI4K2β mRNA-injected embryos also showed the presence of well-differentiated but disorganised notochord as compared to GFP mRNA-injected embryos (Fig.2I&L). Thus, both gain-and loss-of-function of PI4K2β has the effect of disorganising the notochord, similar to other proteins previously implicated in Wnt-PCP pathway e.g. Vangl and Dishevelled ^12, 39^.

As an additional control for the specificity of the phenotypes, we performed rescue experiments by co-injecting morpholinos with full length PI4K2β mRNA. Fig.2Q-R show 500pg of PI4K2β mRNA but not GFP alone (Fig.2O-P) is able to reduce the severity of the observed phenotypes, resulting in a greater percentage of normal and weakly affected embryos i.e. shorter with straight axis. The amount of PI4K2β mRNA for the rescue experiment was limited to 500pg, as higher amounts also resulted in dorsally bent axes (Fig.2J-L). It is likely that this limitation might have prevented the complete rescue of the PI4K2β-depleted embryos.

We also examined effects of depleting embryos of the PI4K2α, both as another control for specificity, and also to compare side-by-side the published phenotypes ^29^ with those for PI4K2β in the same experiment. In agreement with the published data, and in contrast to effects of PI4K2β, embryos injected with 15ng of PI4K2α-Mo into the two dorsal blastomeres at 4-cell stage led to anteriorisation with a shorter axis and an enlarged cement gland (Fig.S2B). Our data is consistent with Pan *et al* 2008 and anteriorisation is thought to be due to suppression of late, posteriorising canonical Wnt signalling. Taken together, our data suggest that PI4K2β normally inhibits or restricts both the canonical Wnt and non-canonical PCP/Wnt pathways.

### PI4K2β differentially affects Wnt signalling pathways

To test molecularly whether PI4K2α and -β were indeed affecting Wnt pathways, we quantified Wnt canonical and non-canonical outputs in embryos depleted for, or overexpressing each PI4K2 isoform using luciferase reporter assays. For canonical Wnt/β-catenin signalling, early embryos were co-injected with Wnt8 mRNA, TOPFLASH β-catenin reporter and CMV-Renilla reporter together with PI4K2α or -β-Mo and/or the corresponding mRNA and assayed at gastrula stage. Fig.3A shows that TOPFLASH-Luciferase reporter activated by Wnt8 was further potentiated by co-injection with PI4K2β-Mo. In contrast, PI4K2α-Mo inhibited Wnt8-induced TOPFLASH-Luciferase reporter activity (Fig.3B), in line with the reported canonical Wnt inhibitory effect ^29^. To control for the specificity, we performed rescue experiments (Fig.3A). Unlike the GFP negative control, full length GFP-PI4K2β mRNA was able to provide substantial luciferase rescue with augmented TOPFLASH activity. Overexpression of PI4K2β by injection of up to 2000pg mRNA did not have significant effect on the Wnt8-induced TOPFLASH-Luciferase reporter activity (Fig.3A) and higher doses were toxic.

**Fig.3.**
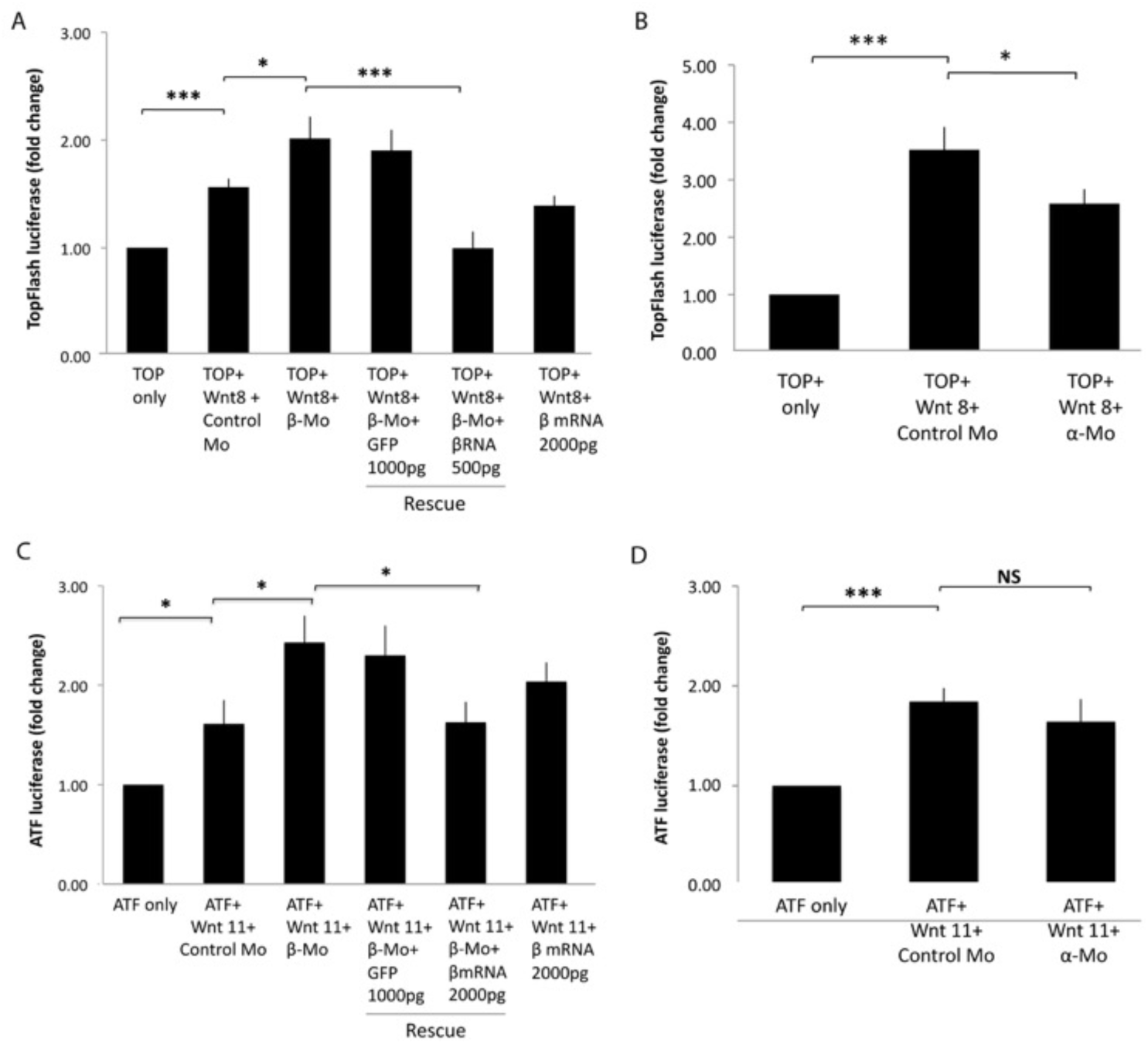
PI4K2β is involved in both the canonical and non-canonical Wnt signalling. (A-D) TOPFLASH/ATF-2-luciferase reporter assays in *Xenopus* embryos show potentiation of Wnt signals by PI4K2β-Mo and rescue by PI4K2β mRNA in contrast to effects of PI4K2α. Embryos were injected at the animal region of two dorsal blastomeres at 4-cell stage with reporter DNAs, morpholinos (Mos) and/or mRNAs as indicated and assayed at stage 11 in pools of 3. Histograms represent three separate samples from each of three independent egg batches for each condition. Values are given as mean ± SEM (Student’s *t*-test without ANOVA ^58^, P < 0.05 *, P < 0.01 **, P < 0.001***).

To determine effects of perturbations on the non-canonical PCP/Wnt signalling, embryos were co-injected with Wnt11 mRNA, ATF-luciferase reporter ^40^ and CMV-Renilla reporter together with PI4K2α or -β-Mo and/or the corresponding mRNA. Fig.3C shows that ATF-luciferase reporter activated by 600pg of wnt11 was further potentiated by co-injection with PI4K2β-Mo, but not PI4K2α-Mo (Fig.3D). To control for the specificity, we performed luciferase rescue experiments. Fig.3C shows that GFP alone did not repress the ATF-luciferase reporter activity, while a substantial luciferase rescue with augmented ATF-luciferase activity using 2000g of full length PI4K2β mRNA. Overexpression of PI4K2β by injection of up to 2000pg mRNA did not have significant effect on the ATF-luciferase reporter activity. It is noted that PI4K2β cannot activate either reporter on its own (data not shown). Taken together, the reporter gene assays confirm that PI4K2β indeed acts on the Wnt8-induced canonical and Wnt11-induced non-canonical PCP signalling pathways.

### PI4K2β is required for Wnt signalling in the receiving tissue

Both PI4K2α/-β have been implicated in Golgi function and protein secretion ^23^ and hence, we wanted to test if PI4K2α/-β-Mo acts on secretion of Wnt or on the response to Wnt. We carried out the animal cap explant sandwich assay, an established method in *Xenopus* for examining cell-autonomous versus non-autonomous effects on signalling ^41, 42^. As described in Materials and Methods and schematized in Fig.4A-C, two groups of embryos (i.e. “sending” versus “receiving”) were injected separately at 2-4-cell stage with Wnt8 mRNA and reporter DNA respectively (Fig.4A). PI4K2α/-β-Morpholinos were co-injected in some embryos with the Wnt8 mRNA to test for effects of PI4K2α/-β depletion on the production or secretion of Wnt, and in some embryos injected with the TOPFLASH reporter to test effects on the response to Wnt. Animal caps were subsequently cut out at stage 8 - 10 and the caps of the sending embryos were conjugated with caps from receiving embryos to form “sandwiches” in which only one contained Mo (Fig. 4B,C). The explants were cultured until they reached stage 12 and harvested for luciferase assay. Fig. 4D,E shows reporter activity induced by Wnt8 was potentiated by co-injection with PI4K2β-Mo and inhibited by PI4K2α-Mo respectively in the whole embryo control or when co-injected in the same half of the conjugate as the reporter (Fig.4D). No significant difference was observed when PI4K2β-Mo was co-injected in the Wnt-overexpressing tissue. Similarly, PI4K2α-Mo in the same experiments inhibited canonical Wnt signal response and had no effect in the Wnt overexpressing tissue (Fig.4E), consistent with its published role in Wnt signal transduction ^29^. These results indicate that both PI4K2α/-β act on the response to a Wnt signal rather than Wnt production or secretion. Unlike PI4K2α, PI4K2β depletion potentiates the canonical Wnt signal response, suggesting that its normal function is inhibitory for this pathway.

**Fig.4.**
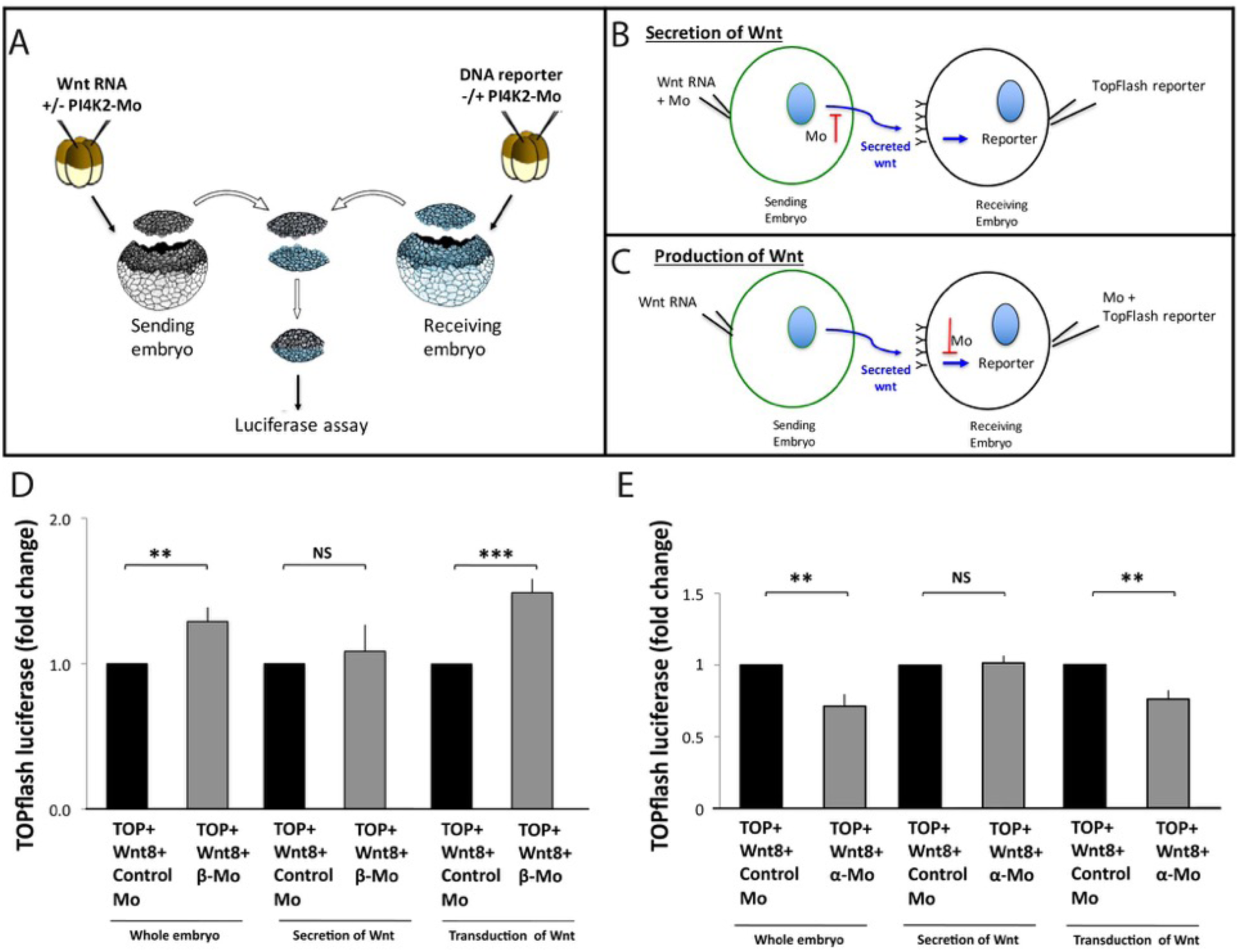
Both PI4K2α/-β-Mo act on signal reception/transduction of Wnt. TOPFLASH luciferase reporter assay in *Xenopus* embryos and explants. A) Schematic diagram of the experimental procedure. See text (Materials & Methods and Results) for details. D&E) Both PI4K2α/-β-Mos potentiated Wnt8-induced TOPFLASH-Luciferase reporter activity when co-injected in the same half of the conjugate as the reporter. Histograms represent luciferase assays from three independent experiments (total of at least 3 pools of 5 sandwiches each per condition per experiment). Values are shown as mean ± SEM

### Phosphorylation of serine-265 by PAR-1 regulates the subcellular localisation of PI4K2β *in vivo*

The primary sequence of PI4K2β contains a PAR-1 phosphorylation consensus site KVGS ^43^, but it has not previously been identified as a PAR-1 substrate. To test whether this site is phosphorylated by PAR-1, we mutated serine (Ser265) to alanine to block PAR-1 phosphorylation. The PI4K2β wild type and the mutated (PI4K2βSA) proteins were transcribed and translated *in vitro* and then incubated with purified GST-tagged human MARK3 wild type (WT) or kinase dead (KD) proteins and analyzed by SDS-PAGE and Western blotting. Fig.5A shows the Ser265Ala mutation abolishes the appearance of a down-shifted band seen with wild type protein confirming Ser265 as a phosphorylation site *in vitro*.

**Fig.5.**
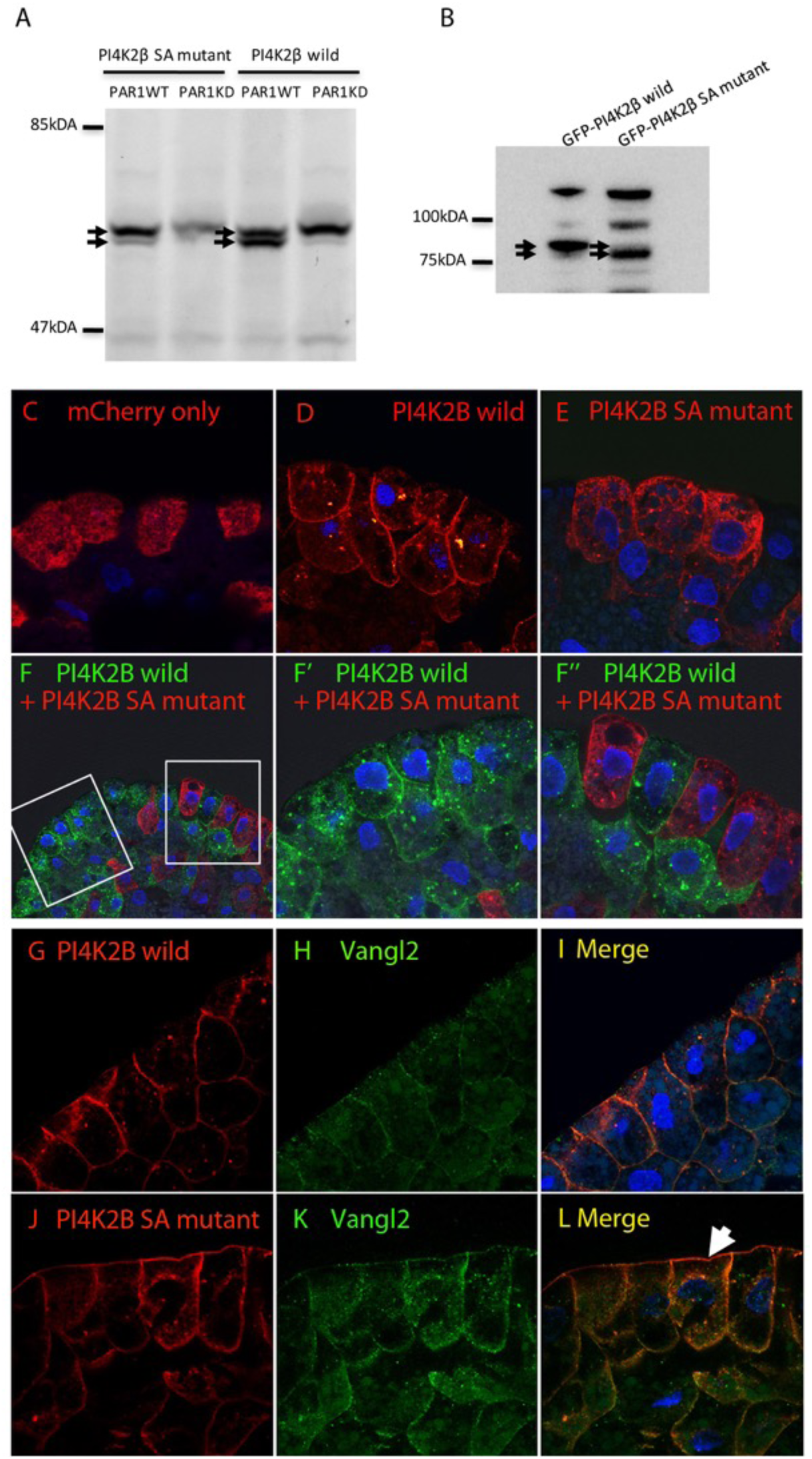
The consensus PAR-1 (Mark2/3) target site on PI4K2β is phosphorylated in vitro and used in vivo to regulate its apicobasal localisation. **(A)** Western blot showing *in vitro*-synthesised PI4K2β shows PAR-1 kinase activity-dependent downshift (strong lower band of doublet) which is lost upon mutation of Ser-265 to Ala. **(B)** Western blot of GFP-tagged and probed PI4K2β wild type or SA mutant showing altered mobility of the latter. **(C-F”)** Confocal images of cryosections of stage 10.5-11 embryonic ectoderm stained using anti-mCherry/GFP primary antibodies to detect mCherry- or GFP-tagged PI4K2α/-β wild type or -SA mutant from injected mRNA encoding the indicated proteins (F’ and F” are boxed details of panel F). mCherry-PI4K2β wild type has a more cell cortical localisation, predominantly in the basolateral cortex of superficial (apicobasally polarised) cells, while the mCherry-PI4K2β SA mutant is seen at both apical and basolateral cell cortex domains. **(G-L)** Confocal images of sections from mCherry-PI4K2β wild type or –SA mutant mRNA-injected embryos as indicated co-stained using antibodies to endogenous Vangl2 protein. Both PI4K2β wild type and SA mutant partially co-localise with endogenous Vangl2 in the superficial ectoderm, ectopically apical in the SA mutant (white arrowhead in panel L). At least 5 embryos per experiment were examined and (C-P) are representative sections of at least three independent experiments.

To test whether this site is also phosphorylated *in vivo*, mRNA encoding GFP-PI4K2β wild type or SA mutant were injected at the animal region of two dorsal blastomeres at 4-cell stage and the mobility of their protein products compared by Western blot analyses. Fig.5B shows that PI4K2 *β* wild type migrated with a reduced, rather than increased, mobility upon gel electrophoresis as compared to PI4K2 *β* SA mutant. These results suggest that serine residue serine-265 within the PAR-1 consensus phosphorylation site is a target for *in vivo* modification, very likely phosphorylation by PAR-1.

To investigate the significance of this site, we studied the subcellular distribution of mCherry-PI4K2β wild type or SA mutant protein and its mCherry-PI4K2α counterparts in *Xenopus* embryonic ectodermal cells and determine if such distribution is affected by PAR-1 phosphorylation (Fig.5C-F). Fig.5C shows overexpression of mCherry alone resulted in a diffuse pattern in the cytoplasm. Similarly, mCherry-PI4K2α wild type and -SA mutant were localised relatively diffusely (Fig.S4A-B). In contrast, mCherry-PI4K2β wild type (Fig.5D) has a striking enrichment in the basolateral cortex of superficial ectoderm cells, and appears to be excluded from the apical domain. It should be noted that this domain in *Xenopus* embryos contains a number of proteins associated with adherens junctions in more mature mammalian epithelia e.g. occludin ^44^. The mCherry-PI4K2β SA mutant (Fig.5E) is localised to both apical and basolateral membranes and appears somewhat less restricted to the cortex compared to the deeper cytoplasm (allowing for the fact that this is also the location of large, impermeable yolk platelets). Co-injection of the two constructs interfered with expression of both, a phenomenon we have observed with other proteins ^45^. However, injection of GFP-PI4K2β wild type and mCherry-PI4K2β SA mutant into nearby parts of same embryo highlighted that GFP-PI4K2β wild type is more basolaterally localised than the mCherry-PI4K2β SA mutant (Fig.5F-F”). Our data suggest that phosphorylation of Serine-265 of PI4K2β by PAR-1 regulates the subcellular localisation of PI4K2β, which is consistent with PAR-1’s larger role in apicobasal polarization in epithelia.

Finally, we asked whether PAR-1-phosphorylation-dependent localisation of PI4K2β might begin to explain its interaction with Wnt-PCP signalling, since the latter has long been known to occur exclusively at or near the apical domain (mostly peri-apically near adherens junctions) and depends on that localisation ^46^. We examined the effect of expression of mCherry-PI4K2β wild type or SA mutant on the localisation of a core PCP protein, Vangl2 (Fig.5G-P). We found that, Vangl2 was co-localised with PI4K2β, being basolateral in wild-type tissue (Fig.5G-I), but apically mislocalised in PI4K2β SA mutant-injected embryos (Fig.5J-P).

## DISCUSSION

In this paper, we have described an *in vitro* screen for novel targets of PAR-1 in which we identified phosphatidylinositol 4-kinase 2-beta (PI4K2β) as one such target (Fig.1). We have further shown that depletion of PI4K2β in *Xenopus* embryos leads to phenotypes characteristic of perturbation of Wnt pathways, particularly the non-canonical PCP pathway required for normal convergent extension (Fig.2). We go on to demonstrate that these Wnt pathways are indeed perturbed by depletion of PI4K2β and rescued by re-expression (Fig.3). It is surprising that both a gain- and loss-of-function of PI4K2β cause similar dose-dependent defects in dorsoanterior structure associated with Wnt signalling – a severely reduced head and a dorsally bent axis. Severe reduction of heads can be due to early loss of canonical Wnt signalling, which in turn leads to reduced organiser size or activity and such “ventralisation” usually has a strong effect in reducing or even abolishing notochord tissue ^47^. This is however not the case since our results show that PI4K2β-depleted embryos have normally differentiated but disorganised notochords. Head reduction can also be due to excess canonical Wnt signalling after midblastula stages or later in development ^48^. Our reporter assays show that PI4K2β is indeed involved in the Wnt8-induced canonical pathway, with depletion producing a modest increase. This suggests an inhibiting role for the endogenous protein and a late excess of canonical Wnt signalling as the cause of the head reduction phenotype. Although PI4K2β has previously been identified as a promoter of canonical Wnt signalling in zebrafish ^49^, it is possible that the regulation of Wnt signalling by PI4K2β is context-dependent as we have yet to demonstrate that xPI4K2β itself is able to induce double axis formation in *Xenopus* embryos (internal communication), a positive functional readout for Wnt activity^50, 51^.

Dorsally bent axes suggest a defect in the control of convergent extension movement, which is regulated by non-canonical Wnt-planar cell polarity ^38^ or Wnt-calcium signalling pathways ^8, 10, 52^. It is well established that the bent-axis phenotype in *Xenopus* is caused by both gain-and loss-of-function of PCP components ^12, 39^. Further link between PI4K2β and non-canonical Wnt pathway is demonstrated by the thickened and disorganised notochord of embryos depleted for, or overexpressing with PI4K2β, suggesting the PCP pathway is indeed disrupted. This is consistent with our reporter assays which show that PI4K2β, in contrast to PI4K2α, inhibits the Wnt11-induced non-canonical Wnt pathway. In addition, our experiments using the animal cap “sandwich” assay show that this inhibitory activity is in Wnt-responding tissue, not in Wnt production or secretion (Fig.4). Taken together, our data suggest that PI4K2β is a functional inhibitor in both canonical and non-canonical Wnt pathways in early *Xenopus* development.

### Par1 phosphorylation excludes PI4K2β from the apical cortex

It is known that PI4K2β contain a PAR-1 phosphorylation consensus site KVGS, but it has yet been identified as a PAR-1 substrate. We mutated serine (Ser-265) of PI4K2β to alanine and the electrophoretic mobility shift assay showed that the Ser-265Ala mutation abolishes the appearance of a down-shifted band seen with wild type protein, confirming that this site is critical for PAR-1 phosphorylation in *in vitro* system (Fig.5). We showed that PI4K2 *β* wild type migrated with a reduced mobility upon gel electrophoresis as compared to PI4K2 *β* SA mutant, suggesting that serine residue 265 is a target for *in vivo* PAR-1 phosphorylation. In addition, PI4K2β wild type and its non-phoshorylatable SA mutant have a more prominent plasma membrane localisation as compared to PI4K2α. We show that PI4K2β wild type is found predominantly in the basolateral cortex of superficial ectoderm cells, while mutation of the PAR-1 phosphorylation site is sufficient to abrogate this localisation. PAR-1 is known to organise other downstream polarity proteins via mutual phosphorylation reactions to exclude each other from their respective regions of the cortex ^53^. For instance, Par1 directly phosphorylates Par3 and exclude it from the apical cortex ^54^. The same mechanism may apply to exclude PI4K2β from cortical regions that contain Par1. We believe that segregation of PI4K2β into basolateral domain of the cell cortex by PAR-1 phosphorylation is essential to its biological functions.

### PAR-1, PI4K2β and non-canonical Wnt signalling

Previously, we showed that different PAR-1 isoforms control different branches of the Wnt signalling pathways ^6^ i.e. Par1A and Par1BX control the canonical branch while PAR-1BY controls the non-canonical Wnt planar cell polarity (PCP) signalling branch. Here, we show how the polarity information of the cell could affect the non-canonical Wnt-PCP pathway via PI4K2β. A key feature of the non-canonical Wnt-PCP pathway is the asymmetric distribution of their core PCP components such as Dishevelled (Dvl) ^30^, Lethal Giant Larvae (Lgl) ^55^ and Van Gogh-like (Vangl) ^5^. Specifically, we showed PI4K2β colocalised with the Vangl2 at the basolateral domain of apicobasally polarised superficial cells in *Xenopus* ectoderm (Fig.5). Vangl has been shown to be involved in the non-canonical PCP pathway via the activation of the Jun N-terminal kinase (JNK) cascade ^18^. Furthermore, dorsal injection of PI4K2β disturbs convergent extension movements and the resulting phenotypes strongly resemble those produced by perturbation of Vangl ^18^, suggesting that these genes may act in a common pathway.

We speculate that the correct localisation of PI4K2β is a prerequisite for the proper function of PCP proteins. This is consistent with the data of Hammond, who reported the function of plasma membrane PI4P is required to contribute to the pool of polyanionic lipids for membrane targeting of proteins ^56^. We believe that the co-localisation of PI4K2β and Vangl2 may involve Dishevelled (Dvl) since Dvl was found to interact with both PI4K2β ^30^ and Vangl in mammalian cells ^57^.

In conclusion, using gain-of-function and loss-of-function approaches, we demonstrate that PI4K2β regulates the vertebrate Wnt-PCP signalling pathway. Our results provide compelling evidence that the novel interaction between PI4K2β and the core PCP component Vang2 links it to the apicobasal polarity regulator PAR-1. These findings represent an important advancement in our understanding of the molecular mechanisms that integrate the establishment of cell polarity with Wnt signalling pathways.

## METHODS

### Screen for novel phosphorylation targets of PAR-1

The full-length, normalized *Xenopus laevis* oocyte cDNA library used in this study was gifted by Vincenzo Costanzo (CRUK, now at IFOM, Milan). An expression library screening method described ^33^ was adapted with modifications. FluoroTect™ Green Lys *in vitro* Translation Labelling System (Promega) was used for protein labelling in place of ^35^ S-methionine in the original protocol. GST-tagged human MARK3 wild type (WT) and kinase dead (KD) proteins were expressed from cDNA clones in pGEX6P-2 vector (gift of Dario Alessi, MRC Protein Phosphorylation Unit, Dundee, UK) and purified as published ^34^. *In vitro-*translated protein pools were subjected to *in vitro* phosphorylation reaction and separated by SDS-PAGE. Gels were rinsed with water and scanned by Typhoon 9210 Imager (GE Healthcare, Amersham, UK) using the green laser (532 nm), emission filter “526 SP Fluorescein, AlexaFluor488”, sensitivity “High” and pixel size at 100 μm. Image files were analysed using ImageJ (http://rsbweb.nih.gov/ij/index.html). For further analysis, full-length *Xenopus laevis* phosphatidylinositol 4-kinase type 2-alpha and -beta (PI4K2α/-β) clones in pExpress-1 vector were purchased from Source Bioscience and were sub-cloned into pCS2plus with eGFP/mCherry at the N-terminus, respectively.

### Ethical Approval

Animal work was approved by the King’s College Institutional Ethics Committee and authorised under a UK Home Office Project Licence to J.B.A.G.

### Preparation of synthetic RNA and microinjection of embryos

Plasmids were linearised using *Not*1 and capped mRNAs were synthesized by *in vitro* transcription using the mMessage mMachine SP6 transcription kit (Life Technologies). The quality of synthesised mRNA was checked by 1% agarose gel electrophoresis. Embryos were obtained by artificial fertilisation, dejellied in 2% cysteine hydrochloride (pH 7.8-8.0) and cultured in 1/10 NAM. Microinjections were carried out in 3% Ficoll at 2- to 4-cell stage in a total volume of 10nl. Amounts injected indicated in the text are per embryo.

### TOPFLASH/ATF Luciferase reporter assay

Embryos were injected at the animal region of the two dorsal blastomeres at 4-cell stage. 50pg TOPFLASH (Upstate), 25pg CMV-Renilla-Luciferase reporter and 6pg Wnt8 mRNA were used for the canonical pathway. 100pg ATF-luciferase reporter ^40^, 0.5pg CMV-Renilla-Luciferase reporter and 600pg Wnt11 mRNA were used for the non-canonical pathway. The reporter plasmids were injected alone or in combination with 15ng PI4K2α/-β-Mo or 500-1500pg PI4K2α/-β mRNAs. Embryos were collected at stage 11 in pools of 3 embryos and assayed for luciferase activity in triplicate (total of >27 injected embryos). Luciferase activity was measured in accordance with the manufacturer’s recommendations (Promega) and 25μl of cell lysate was used for Luciferase detection. The measured luciferase activity was normalised against the measured Renilla reporter reading.

### Animal cap explant conjugation assay

Embryos were injected at the animal region of the two dorsal blastomeres at 4-cell stage with either 6pg Wnt8 mRNA or 50 pg TOPFLASH reporter plus 25 pg Renilla reporter DNAs. To test effects on the response to Wnt, 15 ng PI4K2α or -β-Morpholinos (Mo) were co-injected with the reporter, while to test effects on the production/secretion of Wnt, the Mo was co-injected with the Wnt8 mRNA. Animal caps were excised at mid-blastula (stage 8-9) and Wnt8-injected caps placed against reporter DNA-injected caps for culturing overnight in 3/4NAM at 14°C to early neurula (stage 14) when extracts were made from pools of 5 conjugates and luciferase assays conducted as above.

### Immunofluorescence and hematoxylin staining

Embryos were fixed at stage 10.75 to 12 in MEMFA for 2 hours at room temperature or Dent’s fixative ^6^ for 6 hours at −20°C (rehydrate with gradual change of methanol/PBS) and embedded in fish gelatin (16.7ml of 40% gelatin (Sigma-Aldrich) and 7.5g sucrose in 50ml ddH_2_O) at 4°C overnight. Embedded samples were sectioned using a Cryostat and serial 15μm frozen transverse sections of the entire embryos were collected. For hematoxylin staining, slides were stained with 50% hematoxylin solution (Sigma-Aldrich), washed with tap water, and differentiated with 0.1% HCl. For immunofluorescence, sections were permeabilised and blocked in PBS containing 0.01% Triton X-100 and 5% Goat/Donkey serum at room temperature for 10 minutes. Mouse monoclonal to GFP (clone 3E6, Molecular Probes) 1:500, rabbit/mouse polyclonal to mCherry (Abcam) 1:500, goat polyclonal to Vang2 (N-13, Santa Cruz) 1:250 were used overnight at 4°C. This is followed by Alexi Fluor-488 and/or −568 (Molecular Probes) 1:500 for 1 hour and DAPI 1:10 000 for 5 minutes at room temperature. Samples were mounted with glycerol and analysed with confocal microscope. Confocal images were taken using either a 20×, 40x oil or 63x glycerol objective. For phenotypic studies, embryos were grown to stage 33 to 35 and fixed with 4% PFA. Embryos were scored as normal (white), shorter (light gray), bent (dark gray) and super bent (black) as shown in the Fig.S3.

### Western Blot

Two animal blastomeres of four-cell stage were injected with 500pg eGFP-PI4K2α/-β mRNA. Five injected embryos per sample were lysed at stages 10.5–11.5 in 50ul lysis buffer (1% Triton X-100, 50 mM Tris [pH 7.6], 50 mM NaCl, 1 mM EDTA, 0.1 mM PMSF, 10 mM NaF, 1 mM sodium orthovanadate, 25 mM β-glycerophosphate, Protease Inhibitor Cocktail Complete [Roche]), and the supernatants were centrifuged at 13,000 × g for 10 min. One-third to one embryo equivalents were loaded at per lane on 4%-12% or 8% Bolt® Bis-Tris (Invitrogen). GFP Rabbit IgG Polyclonal Antibody (Molecular Probes) 1:500 and HRP-conjugated goat anti-rabbit IgG (Thermo Scientific) 1:3000 were used sequentially in blocking buffer (1×PBS, 0.1% Tween, 5% BSA).

### Morpholino antisense oligonucleotide injection

Morpholinos (Gene Tools) were re-suspended in water to a concentration of 1mM and stored at-20°C. The following gene specific morpholinos were used in this study: PI4K2α-Mo(ATACTAACGGGCTGGTCTCATCCAT)^29^, PI4K2β*-*Mo-1 (GCTCAGTCGTTTGCTTCTGCTCCAT) and PI4K2β*-*Mo-2 (CTCCCTT CTATTCCCAATGAAAGCA). 10-20ng per embryo was used.

## ACKNOWLEDGMENTS

We thank H. Steinbeisser and P. Angel (DKFZ-ZMBH Alliance) for ATF-2 luciferase reporter, Vincenzo Costanzo (CRUK, now at IFOM, Milan) for the *Xenopus laevis* oocyte cDNA library and Shane Minogue (UCL) for critically reading the manuscript. This work was supported by a BBSRC research grant BB/J015075/1 to J.B.A. Green.

## Author contributions statement

S.M.T, W.C and J.B.A.G designed the research and analysed data; S.M.T, W.C and J.L performed the experiments; S.M.T and J.B.A.G wrote the manuscript. All authors reviewed the manuscript. The authors declare no conflict of interest.

## Supplementary Figures

**Fig.S1.**
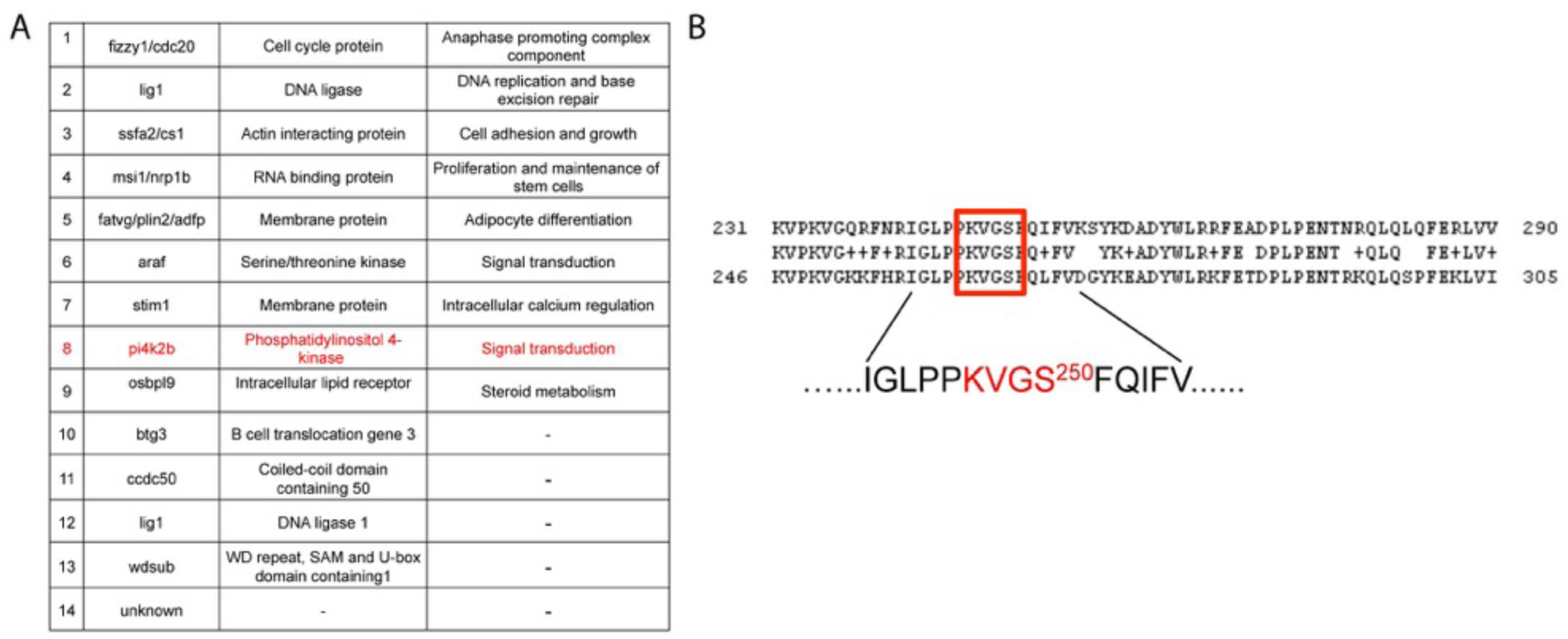
Novel phosphorylation targets of PAR-1. A) The screen identified 14 potential targets with MARK3 phosphorylation and 11 of them were known genes/ homologues. B) Sequence comparison of PAR-1 phosphorylation consensus site KVGS with Xenopus PI4K2α (top) / PI4K2β (bottom). The red box highlights PAR-1 phosphorylation consensus site KVGS.

**Fig.S2.**
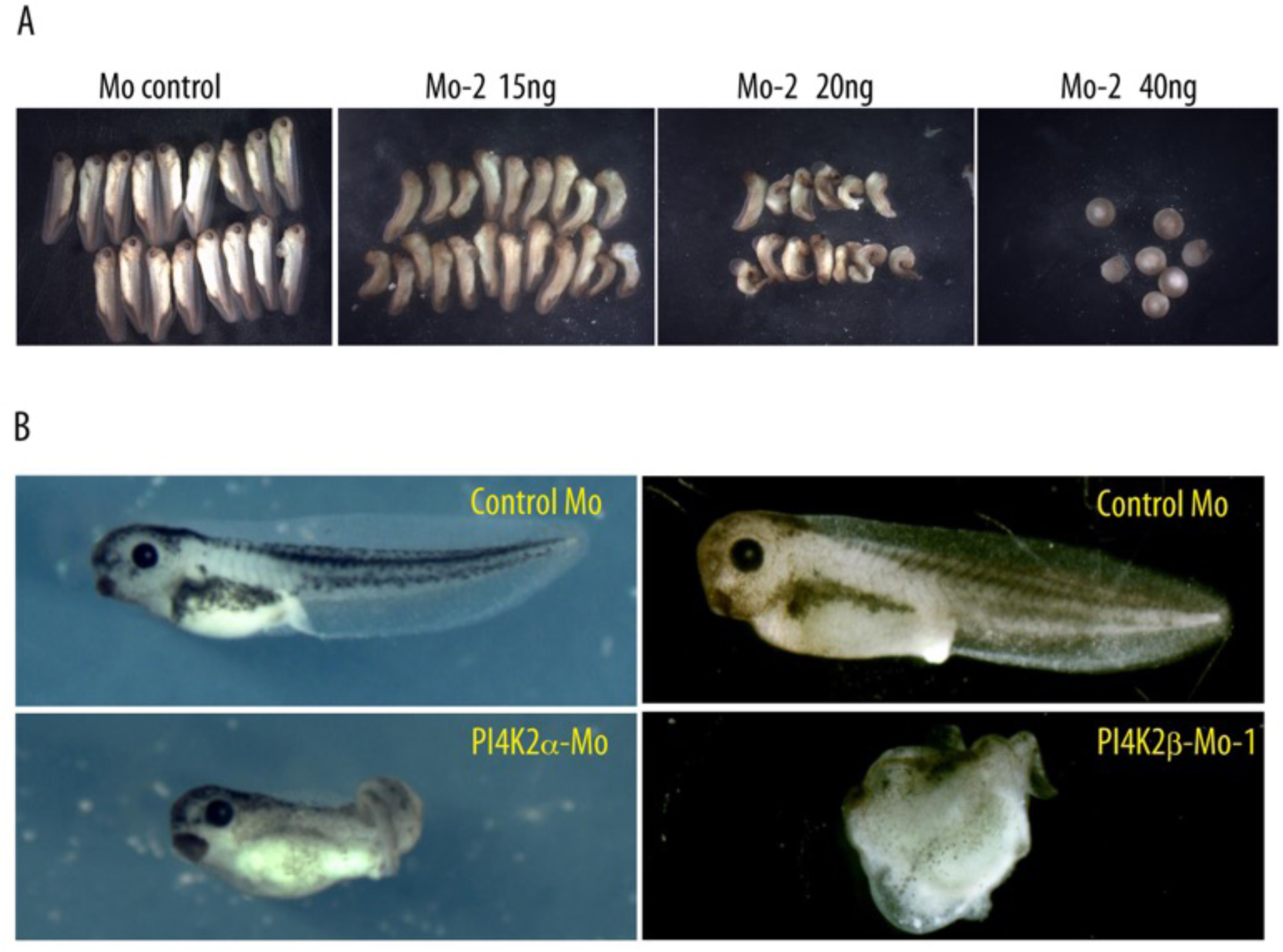
PI4K2α and -β perturbations cause severe but different defects in dorsoanterior structure during embryogenesis. Representative phenotypes at tailbud and tadpole stages of embryos injected with A) PI4K2β-Mo-2 at the doses shown; B) PI4K2α or PI4K2β-Mo-1. Embryos were injected with PI4K2β-Mo-1/2 resulting in distinct dose-dependent defects, whereas PI4K2α-Mo led to anteriorisation with a shorter axis and an enlarged cement gland.

**Fig.S3.**
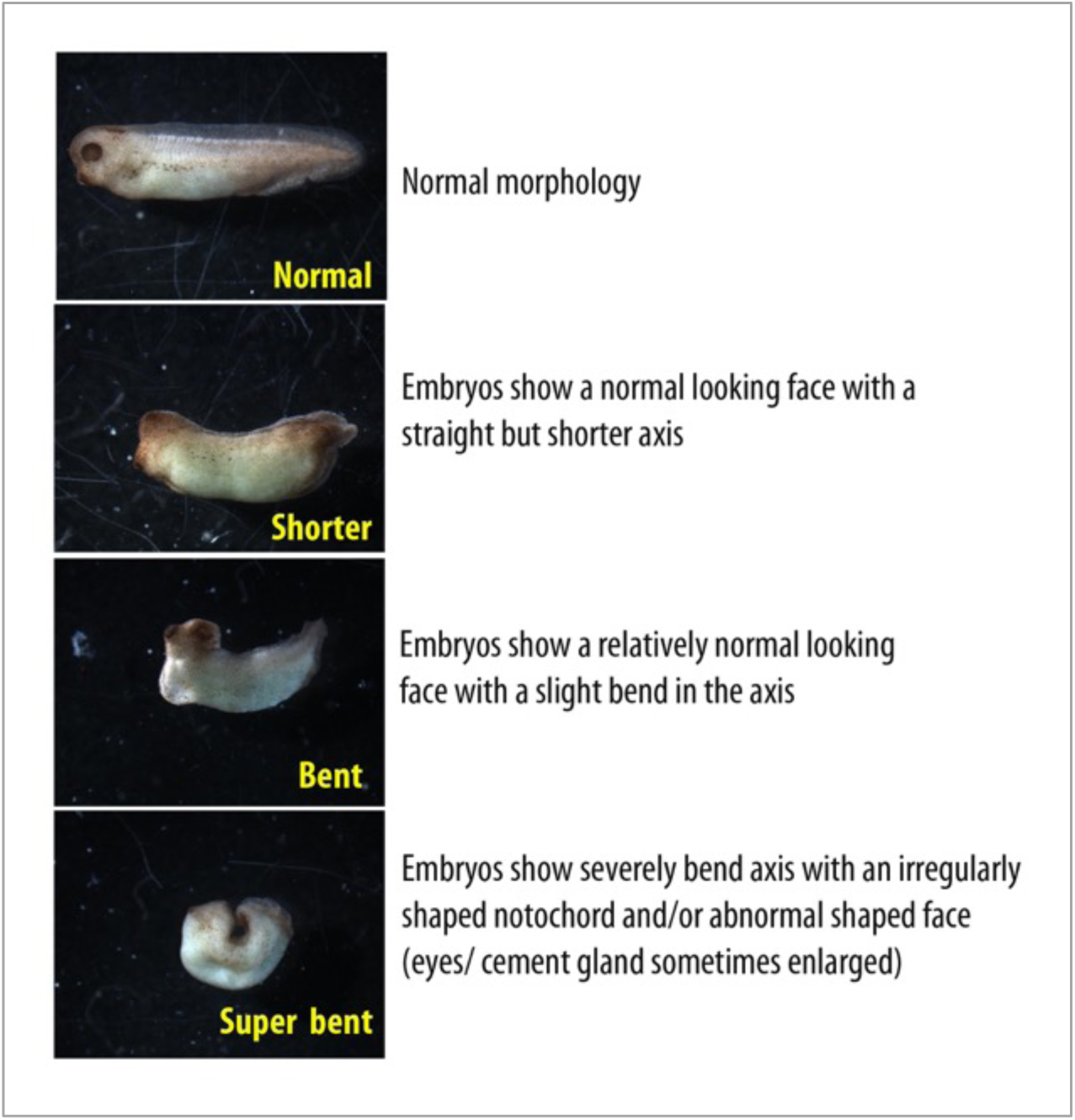
Embryos injected with PI4K2β-Mo cause severe defects in dorsoanterior structure. Representative phenotypes of embryos injected with control Mo 15ng or PI4K2β-Mo 15ng. Embryos were scored at tailbud and tadpole stages as normal (white), shorter (light gray), bent (dark gray) and super bent (black) as shown.

**Fig.S4.**
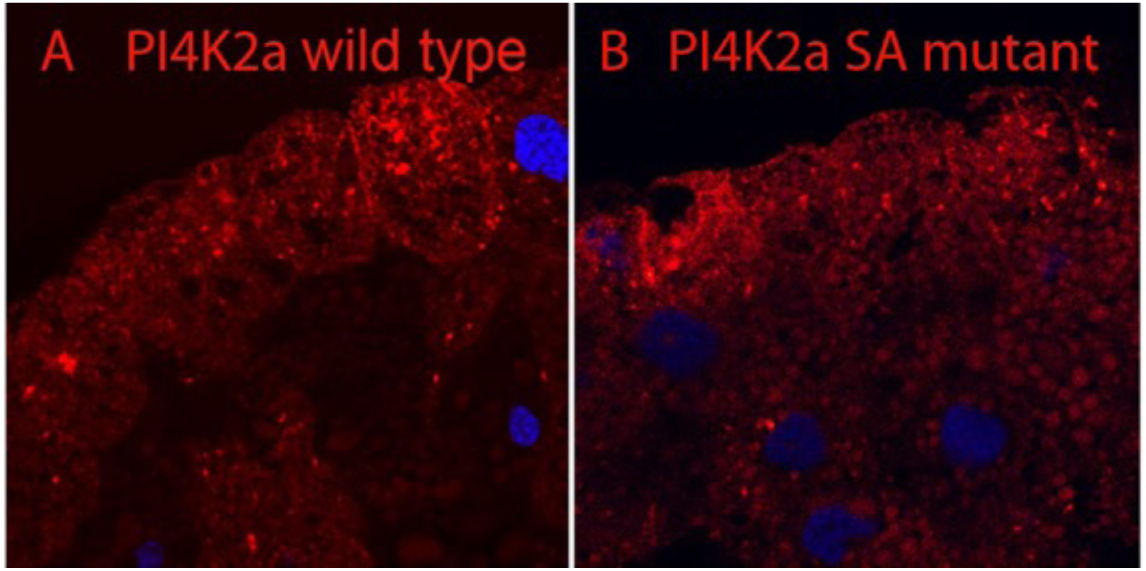
Localisation of PI4K2α wild type or SA mutant in the superficial ectoderm. Embryos were injected with 100pg mCherry-PI4K2α wild type or SA mutant mRNA Cryosections of stage 10.5-11 embryonic ectoderm were stained with anti-mCherry. Both mCherry-PI4K2α wild and SA mutant were localised relatively diffusely.

